# Ventral Pallidum and Amygdala Cooperate to Restrain Reward Approach from Overriding Defensive Behaviors

**DOI:** 10.1101/2023.12.27.573462

**Authors:** Alejandra Hernández-Jaramillo, Elizabeth Illescas-Huerta, Francisco Sotres-Bayón

## Abstract

Foraging decisions involve assessing potential risks and prioritizing food sources, which can be challenging when confronted with changing and conflicting circumstances. A crucial aspect of this decision-making process is the ability to actively suppress defensive reactions to threats (fear) and focus on achieving specific goals. The ventral pallidum (VP) and basolateral amygdala (BLA) are two brain regions that play key roles in regulating behavior motivated by either rewards or threats. However, it is unclear whether these regions are necessary in decision-making processes involving competing motivational drives during conflict. Our aim was to investigate the requirements of the VP and BLA for foraging choices in conflicts involving fear suppression. Here, we used a novel foraging task and pharmacological techniques to inactivate either the VP or BLA, or to disconnect these brain regions before conducting a conflict test. Our findings showed that BLA is necessary for making calculated risky choices during conflicts, whereas VP is necessary for invigorating the drive to obtain food, regardless of the presence of conflict. Importantly, our research revealed that the connection between VP and BLA is critical in limiting risk behaviors when searching for food that requires effort in conflict situations. This study provides a new perspective on the collaborative function of VP and BLA in driving behavior, aimed at achieving goals in the face of danger.

## Introduction

Decisions that have life or death consequences demand a rapid weighing of obstacles and opportunities to take appropriate action in continually changing environments. A particularly intriguing aspect of such survival decision making is the challenge of shifting from secure food acquisition to risk-prone food seeking, which requires active suppression of defensive responses to threats (fear) while pursuing specific goals (Rangel et al., 2008; Headley et al., 2019; Mobbs et al., 2020). When searching for food, foraging animals often face dangers in nature, which illustrates their ability to suppress their fear to achieve their goal. A novel behavioral tool, the crossing-mediated conflict (CMC) task, was recently developed to investigate this choice strategy in rats (Illescas-Huerta et al., 2021). This naturalistic task allows for the assessment of foraging choices that involve rapid transitions from effortful safe food acquisition to food seeking, despite threats within dynamic and unpredictable conditions. In the CMC test, individuals are presented with a foraging challenge that requires them to select an approach based on prior associations they have learned. The test involves crossing an alley to obtain food, and rats must decide whether to cross to obtain food safely or confront their fear and cross despite the danger involved. The relatively understudied ability in this task highlights the necessity of researching the neural processes that govern decisions when there is a struggle between rewards and threats.

The ventral pallidum (VP) and basolateral amygdala (BLA) are critical neural substrates involved in orchestrating motivated behaviors driven by rewards and threats (Smith et al., 2009; Moscarello and LeDoux, 2013; Janak and Tye, 2015; Root et al., 2015; Bernardi and Salzman, 2017). The VP has traditionally been thought of as the motor output of the basal ganglia, responsible for guiding reward-seeking behaviors. However, recent research has called this understanding into question. Emerging research has demonstrated that VP neuronal activity exhibits dynamic responsiveness to relative threats (Moaddab et al., 2021) and mediates avoidance behaviors (Saga et al., 2017; Faget et al., 2018; Stephenson-Jones et al., 2020), positioning VP uniquely to detect and process threat-related information. Additionally, VP has been implicated in encoding expected reward values that influence incentive motivation for choice behaviors (Tachibana and Hikosaka, 2012; Richard et al., 2016; Richard et al., 2018; Fujimoto et al., 2019; Ottenheimer et al., 2020a). Similarly, the conventional notion of BLA in threat-related memory expression has changed over time (Pare and Quirk, 2017), and accumulated evidence has shown that BLA neurons also respond to reward cues (Beyeler et al., 2016). This dual function of VP and BLA highlights their potential roles in decision-making processes shaped by opposing motivational drives. Consistent with this notion, studies employing food-seeking behavior or threat-related responses have shown that VP inactivation reduces motivation for effortful food-seeking behavior (Farrar et al., 2008; Farrell et al., 2021; Lederman et al., 2021), whereas BLA inactivation increases the drive to approach threat-associated cues, potentially influencing riskier foraging decisions (Choi and Kim, 2010; Bravo-Rivera et al., 2014; Burgos-Robles et al., 2017; Orsini et al., 2017). Despite these advancements, the extent to which VP and BLA interact and influence foraging choices when competition occurs remains unknown.

Here, we investigated the individual and collaborative necessity of VP and BLA to influence foraging decisions in conflict situations. Specifically, we aimed to study how these brain regions contribute when animals choose between making minimal effort to safely obtain food and engaging in more demanding actions that involve fear suppression. To investigate this, we pharmacologically inactivated the VP and BLA or disconnected these brain regions before the CMC test. Our findings reveal that BLA is necessary for food-seeking decisions that involve taking risks during conflicts, whereas VP primarily regulates decisions related to the motivational drive to seek food, irrespective of the presence of conflict. Notably, our research underscores the importance of connectivity between VP and BLA in restraining risk behaviors associated with actions to seek food while facing danger.

## Results

Rats were trained in the Crossing-Mediated Conflict (CMC) task to discern between safe (non-conflict) and risky (conflict) crossings in order to obtain food in the straight alley apparatus depicted in **Figure 1A**. Testing involved pseudorandom presentations of non-conflict and conflict trials, without shocks (**Figure 1B**).

**Figure 1.**
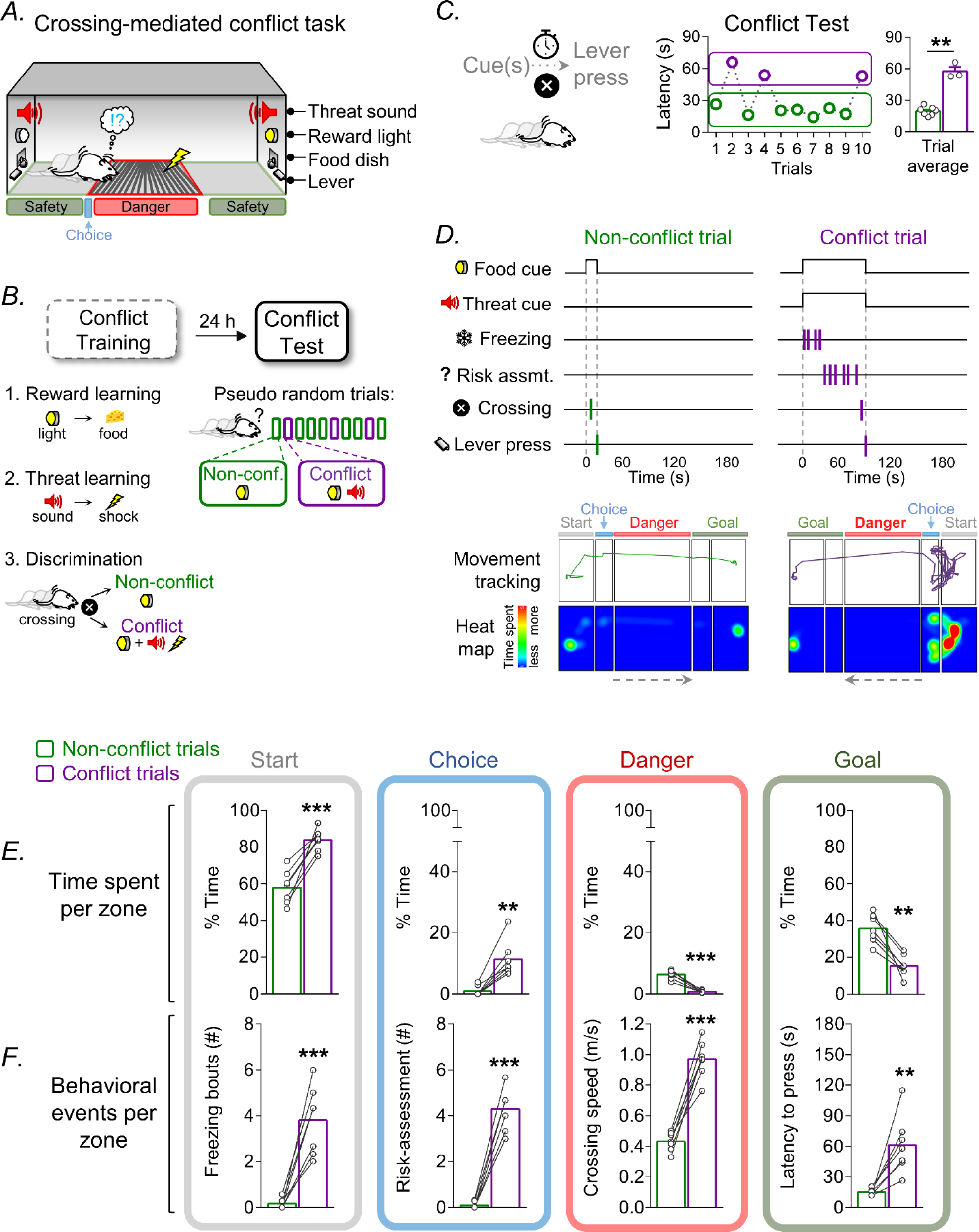
Crossing-mediated conflict task to study the act of overcoming threats for food. **A.** Rats (n=7) were trained and tested in a grid-divided alley foraging between feeding zones, based on cues denoting either conflict or non-conflict situations. **B.** Training involved three consecutive stages: acquisition of reward learning through a light cue in a safety zone, acquisition of threat learning via a white noise cue in a danger zone, and discrimination between cue-guided crossing trials involving conflict (simultaneous light and noise) and those without conflict (light only). Subsequent testing assessed memory-guided decision making during both conflict and non-conflict trials. **C.** Representative rat data indicate prolonged latency in crossing and lever pressing on the opposite side of the alley during conflict trials compared to non-conflict trials within the same session. **D.** Example ethogram, movement tracking, and heat map during the first two consecutive non-conflict and conflict trials showing the relative timing of cues (food and threat), crossing, reward-seeking behavior (lever press), and threat-related responses (freezing and risk assessment). **E.** Analysis of time spent per zone shows that rats spent more time in start and choice zones but less time in danger and goal zones during conflict compared to non-conflict trials. **F.** Analysis of behavioral events per zone shows increased defensive responses and crossing latency during conflict compared to non-conflict trials. **p<0.01 and ***p<0.001 in a Student’s t-test (paired, two-tailed). The error bars in these and subsequent figures represent the SEM, whereas each individual rat is represented by small empty circles.

### Calculated risk-taking incentivized by food-seeking behavior

To examine cue-triggered behavior in the conflict test, we used a group of unimplanted male rats (n=7). **Figure 1C** displays data from a representative rat, showing the extended reaction times (latency from start to goal zone) for crossing to lever-press in conflict trials compared to non-conflict trials (conflict trials: 57.56 s, non-conflict trials: 19.72 s; Student’s two-tailed paired *t*-test, t(2)= 16.65, p= 0.003). **Figure 1D** further reveals the temporal (top) and spatial (bottom) patterns associated with cues, crossing, reward-seeking actions, and responses to potential threats in the same representative rat. Notably, when food-seeking actions coincided with potential threats (conflict trials), a calculated decision-making process emerged, involving both passive (freezing) and active defensive (risk assessment) behaviors, in contrast to non-conflict trials.

Both non-conflict and conflict trials required crossing to access food, with conflict trials involving added complexity, as rats appeared to evaluate the time and effort involved in their decision to cross despite potential danger. Our analysis of time allocation and behavioral events within delimited zones (start, choice, danger, and goal zones) during both conflict and non-conflict trials revealed distinct temporal and spatial profiles (**Figure 1E**). In conflict trials, rats allocated more time to the start and choice zones (start zone, conflict trials: 84.01%, non-conflict trials: 58.07%, Student’s two-tailed unpaired *t*-test, t(6)= 6.96, p< 0.000; choice zone, conflict trials: 11.38%, non-conflict trials: 0.96%, Student’s two-tailed unpaired *t*-test t(6)= 5.58, p = 0.001), while spending less time in the danger and goal zones (danger zone, conflict trials: 0.74%, non-conflict trials: 6.36%, Student’s two-tailed unpaired *t*-test, t(6)= 9.97, p< 0.000; goal zone, conflict trials: 15.24%, non-conflict trials: 35.56%, Student’s two-tailed unpaired *t*-test, t(6)= 5.95, p = 0.001), suggesting a cautious and calculated approach in the face of threats. Furthermore, a comparison of behavioral events showed marked differences between conflict and non-conflict trials (**Figure 1F**). Conflict trials revealed the emergence of freezing events in the start zone (conflict trials: 3.80 events, non-conflict trials: 0.16 events, Student’s two-tailed unpaired *t*-test, t(6)= −6.50, p< 0.000) and the appearance of risk assessment events in the choice zone (conflict trials: 4.28 events, non-conflict trials: 0.08 events, Student’s two-tailed unpaired *t*-test, t(6)= - 13.62, p< 0.000). Rats demonstrated faster crossing of the danger zone (conflict trials: 0.96 m/s, non-conflict trials: 0.46 m/s, Student’s two-tailed unpaired *t*-test, t(6)= −10.72, p< 0.000) and longer latency in pressing the lever in the goal zone (conflict trials: 61.41s, non-conflict trials: 15.09 s, Student’s two-tailed unpaired *t*-test, t(6)= 4.28, p= 0.005) during conflict trials. In summary, our analysis emphasizes the significant temporal investment required to suppress defensive responses in favor of food-seeking actions during conflict trials, representing a calculated risk-taking strategy when confronted with challenging and potentially threatening situations. Thus, choosing food-seeking actions over defensive reactions involves calculating risk during conflict.

### BLA mediates risky food-seeking choices during conflict

To examine the role of the BLA in decision-making between seeking food and defensive reactions in response to threats, we employed pharmacological inactivation of this limbic structure before subjecting the rats to a memory conflict test (**Figure 2A**). Our findings revealed a significant impact of BLA inactivation on the temporal aspects and behavioral events during conflict trials, while no discernible effect was observed during non-conflict trials (**Figure 2B-D**). BLA inactivation during conflict trials rendered all behavioral measures similar to those observed in non-conflict trials. Specifically, during conflict trials, BLA inactivation reduced the time spent in the start zone, aligning with the durations observed in non-conflict trials (factorial ANOVA, group: F(1,22)= 33.98, p= 0.000; trials: F(1,22)= 16.71, p= 0.000; interaction: F(1,22)= 8.71, p= 0.007; conflict trials SAL: 85.52 s; conflict trials INACT: 49.46 s, post hoc comparison p= 0.000; conflict trials INACT vs. non-conflict trials VEH, post hoc comparison p= 0.61, conflict trials VEH vs. non-conflict trials INACTIV, post hoc comparison p= 0.79). This finding suggests that BLA plays a pivotal role in delaying food-seeking decisions when faced with threats, as reflected by the decrease in time spent in the start zone. Moreover, BLA inactivation significantly diminished defensive responses, including freezing in the start zone and risk assessment in the choice zone (freezing bouts, factorial ANOVA, group: F(1,22)= 10.57, p= 0.003; trials: F(1,22)= 14.45, p= 0.000; interaction: F(1,22)= 10.65, p= 0.003; conflict trials VEH: 2.2 events; conflict trials INACTIV: 0.25 events, post hoc comparison p= 0.000; risk assessment, factorial ANOVA, group: F(1,22)= 9.03, p= 0.006: trials: F(1,22)= 13.55, p= 0.001; interaction: F(1,22)= 8.93, p= 0.006; conflict trials VEH: 1.70 events; conflict trials INACT: 0.20 events, post hoc comparison p=0.001). Notably, BLA inactivation also resulted in increased time spent within the danger zone and slower crossing speed (danger zone, factorial ANOVA, group: F(1,22)= 15.42, p= 0.000; trials: F(1,22)= 9.00, p= 0.006; interaction: F(1,22)= 12.39, p= 0.001; conflict trials VEH: 1.40%; conflict trials INACT: 8.31%, post hoc comparison p= 0.000; crossing speed, ANOVA, group: F(1,22)= 2.48, p= 0.12; trials: F(1,22)= 12.74, p= 0.001; interaction: F(1,22)= 19.09, p= 0.000; conflict trials VEH: 0.67 m/s, conflict trials INACT: 0.42 m/s; post hoc comparison, p= 0.002), indicating that the BLA plays a vital role in promoting the rapid crossing of this dangerous area to achieve a goal. Additionally, rats with BLA inactivation exhibited an extended duration of time spent in the goal zone, but a shorter time to press for food (goal zone, factorial ANOVA, group: F(1,22)= 29.46, p= 0.000; trials: F(1,22)= 14.03, p= 0.001; interaction: F(1,22)= 5.61, p= 0.02; conflict trials VEH: 13.06%; conflict trials INACT: 42.21%, post hoc comparison p=0.000; latency to press in the goal zone, factorial ANOVA, group: F(1,22) = 17.82, p= 0.000; trials: F(1,22)= 19.91, p= 0.001; interaction: F(1,22)= 18.11, p= 0.000; conflict trials VEH: 43.07 s; conflict trials INACT: 17.68 s, post hoc comparison p= 0.000). After inactivation of the BLA, the decrease in latency to cross was comparable to that previously observed using the anxiolytic drug diazepam (Illescas-Huerta et al., 2021). Overall, these findings suggest that BLA not only plays a critical role in the cautious decision-making process when confronting threats for food but also signals urgency, promoting the immediate compromise of potential harm to achieve a goal.

**Figure 2.**
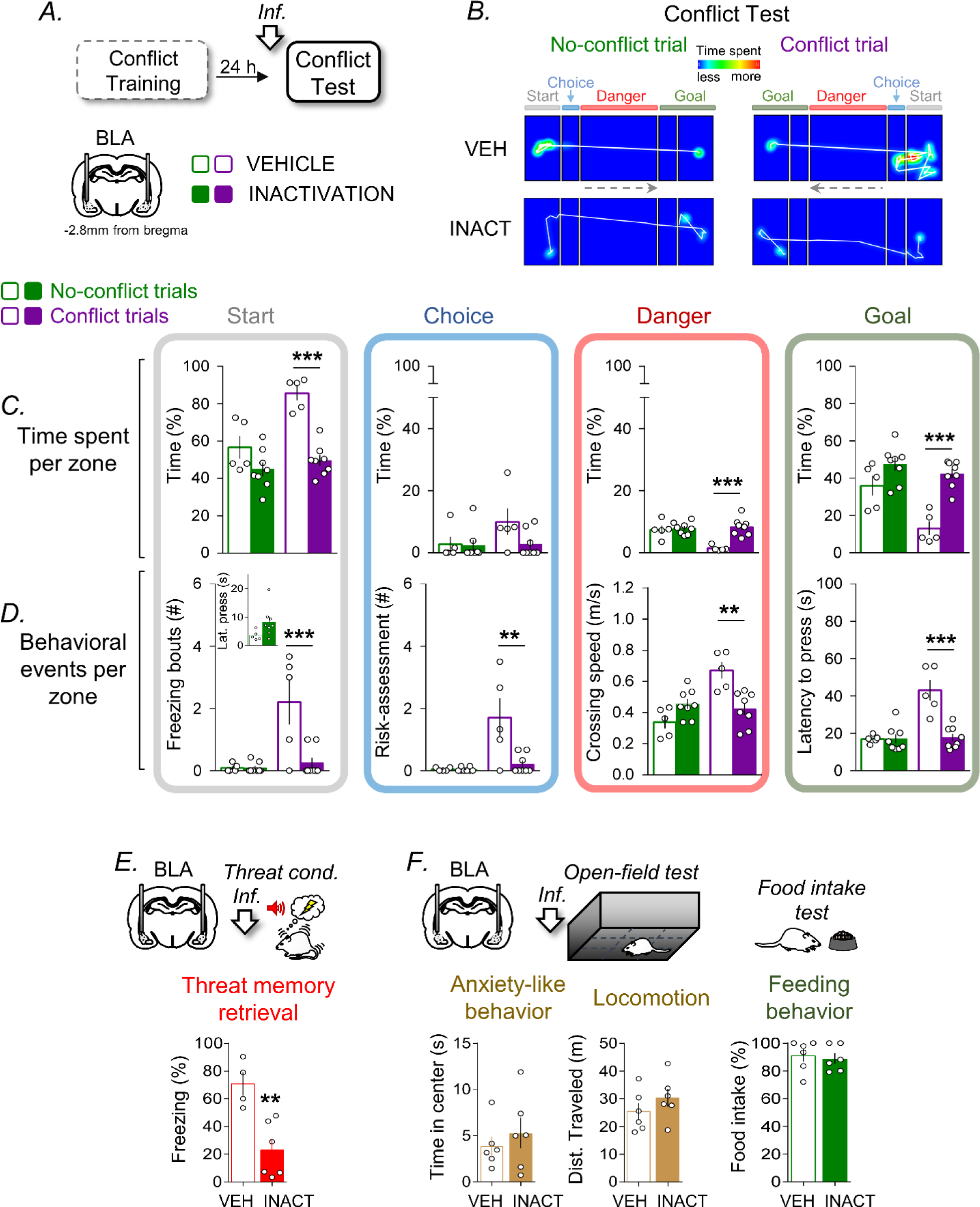
BLA inactivation overcomes the fear to obtain food by preventing conflict-triggered threat-related behaviors. **A.** Rats were infused with vehicle saline solution (VEH, n = 5) or a cocktail of muscimol and baclofen (INACT, n = 8) into the BLA before the conflict test. **B.** Overlaid movement tracking and heat maps of representative rats (VEH and INACT) during the first two consecutive non-conflict and conflict trials of the conflict test. **C.** Analysis of the time spent per zone showed that BLA inactivation decreased the time spent in the start safety zone while increasing it in the danger and goal zones during conflict trials without affecting non-conflict trials. **D.** Analysis of behavioral events per zone shows that BLA inactivation impaired defensive responses (freezing and risk assessment) in the start zone, slowed crossing speed in the danger zone, and decreased crossing latency to press in the goal zone during conflict trials without affecting non-conflict trials. BLA inactivation did not affect latency to press on the same side of the alley (inset). **E.** BLA inactivation impaired the retrieval of auditory threat conditioning memory, as indicated by low levels of freezing in the inactivation group (n=4) compared to the vehicle group (n=6). **F.** BLA inactivation did not affect anxiety-like behavior and general locomotion in the open-field test, as well as not affect feeding behavior in a free food intake test, as indicated by similar time in the center of the open-field and similar levels in the other behavioral measurements comparing vehicle (n=6) and inactivation (n=6) groups. *p<0.05, **p<0.01, and ***p<0.001 in post hoc comparison after ANOVA and Student’s t-test.

Consistent with previous results (Muller et al., 1997; Sierra-Mercado et al., 2011), we found that BLA inactivation impaired threat memory retrieval (**Figure 2E and F**; threat memory, VEH: 70.66%, INACT: 22.97%, Student’s two-tailed unpaired *t*-test, t(8)= 3.91, p= 0.004) while leaving anxiety-like behavior, general locomotion, and consummatory feeding behavior unaffected (anxiety-like behavior, VEH: 3.88 s, INACT: 5.25 s, Student’s two-tailed unpaired *t*-test, t(10)= 0.71, p= 0.49; locomotion, VEH: 25.39 m, INACT: 30.32 m, Student’s two-tailed *t-*test, t(10)= 1.12, p= 0.28; feeding behavior, VEH: 91.15%, INACT: 88.86%, Student’s two-tailed *t-*test, t(10)= 0.38, p= 0.70). Consistent with the lack of effect of BLA inactivation on feeding behavior, cued reward-seeking responses (as depicted in the **inset in Figure 2D**; VEH: 3.54 s, INACT: 8.27s, Student’s two-tailed unpaired *t*-test, t(11)= −1.93, p= 0.078) also remained unaltered. In summary, our results support the notion that BLA inactivation impairs the retrieval of learned threat-related behaviors, thereby allowing the expression of food-seeking behaviors even in the face of potential harm. This suggests that BLA normally inhibits risky behaviors and facilitates a cautious approach when confronted with the challenge of obtaining food. Thus, BLA may restrain the choice of seeking food over defensive reactions during conflict.

### VP mediates the motivation drive for food-seeking, irrespective of conflict

VP plays a pivotal role in incentive motivation involving the invigorating effort to seek food (Farrar et al., 2008; Lederman et al., 2021), which leads to the intriguing possibility that it also mediates the additional challenge that represents risky food seeking. To evaluate the specific involvement of the VP in confronting threats for food, we employed pharmacological inactivation of this ventral basal ganglia structure before the memory conflict test (**Figure 3A-B**). Notably, our findings revealed that VP inactivation significantly reduced the time spent and the occurrence of risk assessment events in the choice zone during conflict trials (choice zone, factorial ANOVA, group: F(1,24)= 18.67, p= 0.000; trials: F(1,24)= 25.23, p= 0.000; interaction: F(1,24)= 18.40, p= 0.000; conflict trials VEH: 14.54%; conflict trials INACT: 1.51%, post hoc comparison, p= 0.000; risk assessment, factorial ANOVA, group: F(1,24)= 6.80, p = 0.015; trials: F(1,24)= 24.51, p= 0.000; interaction: F(1,24)= 7.04, p= 0.013; conflict trials VEH: 6.6 events; conflict trials INACT: 2.09 events, post hoc comparison p= 0.005) (**Figure 3C-D**). This finding suggests that VP is not only critical in incentive motivation for food-seeking but also in the complex process of weighing potential risks and rewards associated with such choices. Moreover, VP inactivation led to a substantial increase in the time required by rats to obtain food, regardless of whether the task involved confronting a threat (as in conflict trials) or not (as in non-conflict trials) (factorial ANOVA, group: F(1,24)= 27.38, p= 0.000; trials: F(1,24)= 6.18, p= 0.02; interaction: F(1,24)= 1.28, p= 0.26; non-conflict trials VEH: 17.04 s, non-conflict trials INACT: 131.58 s, post hoc comparison p= 0.000; conflict trials VEH: 82.17 s, conflict trials INACT: 155.94, post hoc comparison p= 0.03). This VP inactivation effect is comparable to providing free access to food (satiety) one day before the CMC test (Illescas-Huerta et al., 2021), indicating that the task involves goal-directed behavior guided by incentive motivation. Nearly half of the rats subjected to VP inactivation (three out of seven) failed to cross within the allotted time (180s), suggesting a compromised incentive motivation to obtain food, regardless of the trial type. This apparent lack of motivation was also evident in the VP-inactivated rats’ reluctance to press for food within the start zone (without the need to cross), as depicted in the inset of **Figure 3D** (VEH: 3.38 s, INACT: 92.32 s, Student’s two-tailed unpaired *t*-test, t(12) = −2.81, p= 0.015). Thus, these findings support the role of VP in incentive motivation for food seeking and risk evaluation.

**Figure 3.**
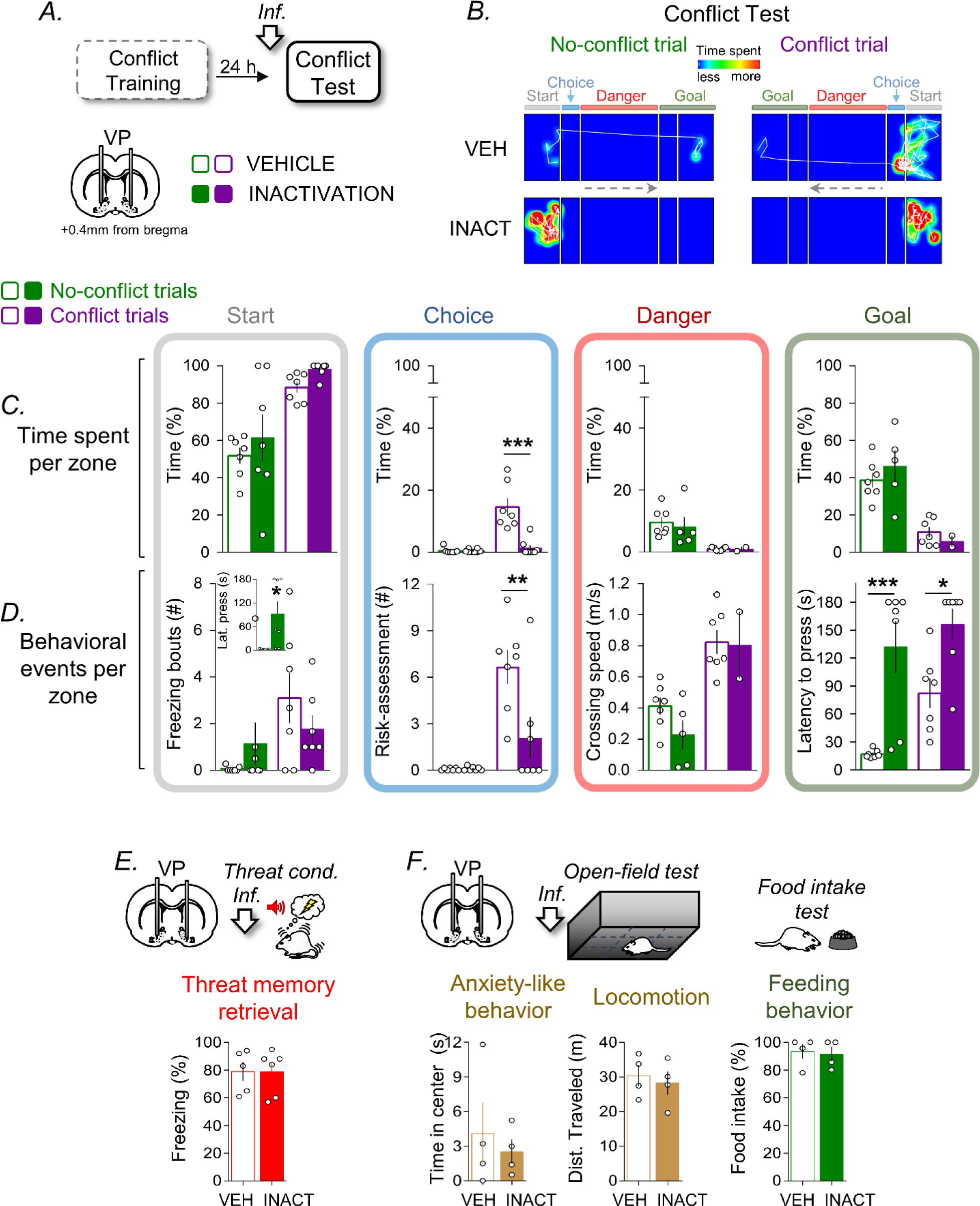
VP inactivation impairs effort-related actions and conflict-triggered risk taking. **A.** Rats were infused with vehicle saline solution (VEH, n = 7) or muscimol (INACT, n = 7) into the VP before the conflict test. **B.** Overlaid movement tracking and heat maps of representative rats (VEH and INACT) during the first two consecutive non-conflict and conflict trials of the conflict test. **C.** Analysis of the time spent per zone showed that VP inactivation decreased the time spent in the start safety zone while increasing it in the danger and goal zones during conflict trials without affecting non-conflict trials. **D.** Analysis of behavioral events per zone shows that VP inactivation decreased risk assessment related to conflict in the choice zone while increasing latency to cross irrespective of trial type. VP inactivation also increased latency to press on the same side of the alley (inset). **E.** VP inactivation did not affect retrieval of auditory threat conditioning memory, as shown by similar freezing levels in the vehicle group (n=5) as compared to the inactivation group (n=6). VP inactivation did not affect anxiety-like behavior and general locomotion in the open-field test, nor did it affect feeding behavior in the free food intake test, as indicated by similar time in the center of the open field and similar levels in the other behavioral measurements comparing vehicle (n=4) and inactivation (n=4) groups. *p<0.05, **p<0.01, and ***p<0.001 in post hoc comparison after ANOVA.

Consistent with previous research, our findings support the idea that VP plays a pivotal role in incentive motivation (Tindell et al., 2004; Richard et al., 2016; Ottenheimer et al., 2018). However, given the observed impairment in risk assessment in the CMC task following VP inactivation and the recent implications of VP in threat-related processing (Peczely et al., 2014; Lenard et al., 2017; Faget et al., 2018; Moaddab et al., 2021), we conducted separate experiments in different rat groups to explore whether VP inactivation is necessary for learning and innate threat-related behaviors (**Figure 3E-F**). The results of these additional experiments indicated that VP inactivation did not influence freezing levels during the retrieval of a threat memory (VEH: 79.0%, INACT: 78.66%, Student’s two-tailed unpaired *t*-test, t(9)= 0.03, p= 0.97), nor did it affect anxiety-like behavior in the open-field test (VEH: 4.13 s, INACT: 2.53 s, Student’s two-tailed unpaired *t*-test, t(6)= 0.56, p= 0.59). These results are consistent with the lack of VP inactivation effect on freezing bouts in the start zone during the CMC test. Furthermore, VP inactivation had no discernible effect on general locomotion and feeding behavior in a free food intake test (distance traveled, VEH: 30.34 m, INACT: 28.24 m, Student’s two-tailed unpaired *t*-test, t(6)= −0.46, p= 0.65; food intake, VEH: 93.46%, INACT: 91.48%, Student’s two-tailed unpaired *t*-test, t(6)= 0.27, p= 0.79). In summary, our findings suggest that VP plays a critical role in the motivational control of goal-directed actions, encompassing those that involve effort, whether associated with rewards or threats.

### VP-BLA connectivity mediates restraint of risky reward choice behaviors

Despite the extensively reported roles of VP and BLA in motivated behaviors and their extensive reciprocal anatomical connections (Zaborszky et al., 1984; Mitrovic and Napier, 1998; Mascagni and McDonald, 2009), the functional significance of the communication between these structures in motivation has been sparsely explored. We showed that BLA inactivation promotes risky food-seeking decisions during conflict, whereas VP inactivation restrains effort-based food-seeking behaviors, regardless of trial type. Consequently, disrupting the exchange of information between the VP and BLA is likely to exert a substantial impact on a rat’s behavior, particularly during conflicts involving food seeking and defensive reactions. To investigate this possibility, we conducted a disconnection experiment by comparing the effects of VP and BLA inactivation on the opposite hemisphere side (contralateral) with their effects on the same hemisphere side (ipsilateral). Given that most communication between brain structures occurs ipsilaterally, contralateral inactivations were employed to disrupt the communication between the VP and BLA (**Figure 4**), whereas ipsilateral inactivations served as controls (**Figure 5**), where communication between structures was presumably minimally disrupted. Thus, ipsilateral inactivation spares VP-BLA communication in one hemisphere, whereas contralateral inactivation disrupts VP-BLA communication in both hemispheres.

**Figure 4.**
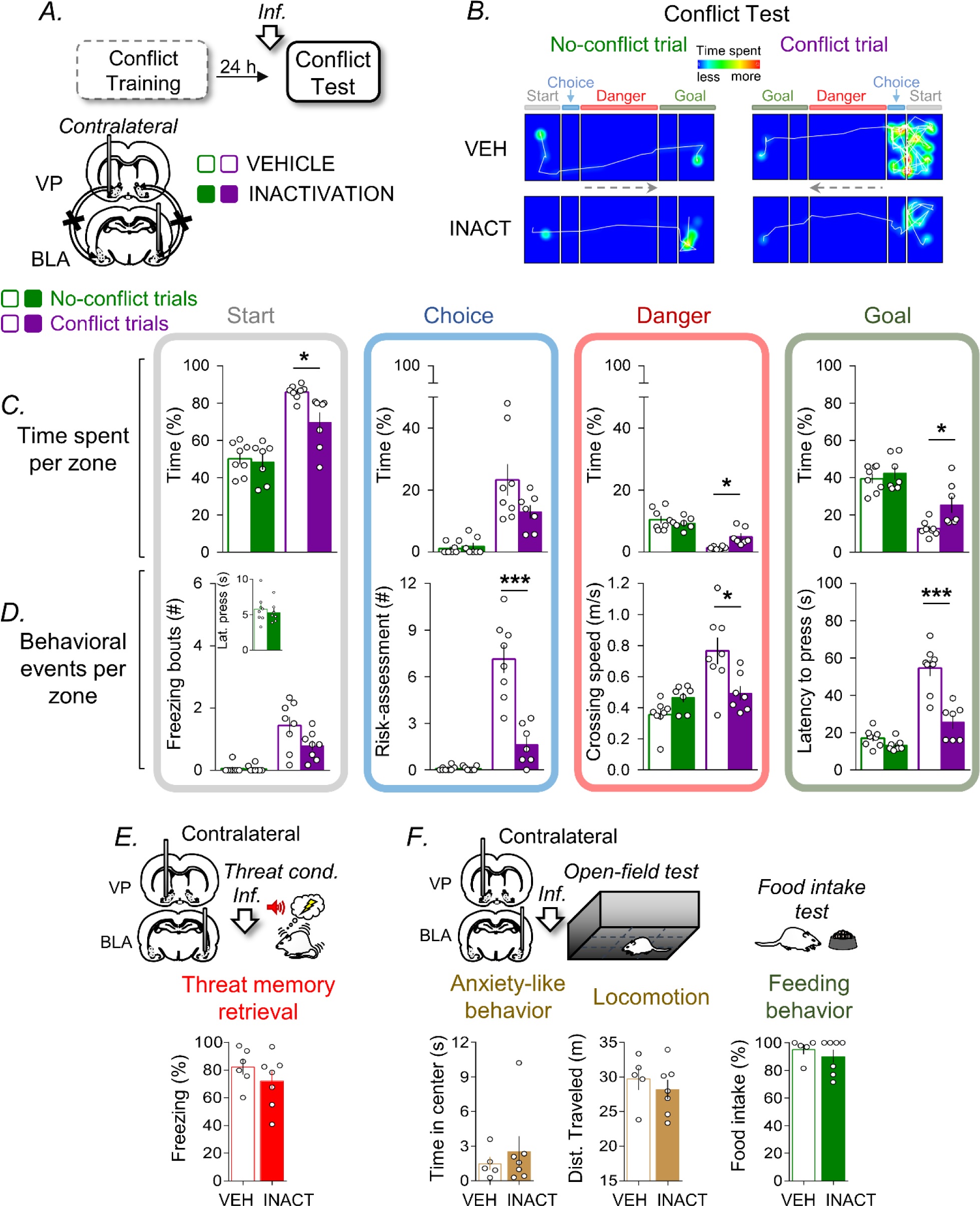
Contralateral VP-BLA inactivation impairs conflict-induced fear in food-seeking efforts. **A.** To functionally disconnect VP-BLA communication, rats were infused into the VP and BLA on the contralateral sides with vehicle saline solution (VEH, n=8) or muscimol into VP and a cocktail of muscimol and baclofen into BLA (INACT, n=7) before the conflict test. **B.** Overlaid movement tracking and heat maps of representative rats (VEH and INACT) during the first two consecutive non-conflict and conflict trials of the conflict test. **C.** Analysis of the time spent per zone showed that VP-BLA contralateral inactivation decreased the time spent in the start zone while increasing it in the danger and goal zones during conflict trials without affecting non-conflict trials. **D.** Analysis of behavioral events per zone shows that VP and BLA contralateral inactivation impaired defensive responses (freezing and risk assessment) in the start zone, slowed crossing speed in the danger zone, and decreased crossing latency to press in the goal zone during conflict trials without affecting non-conflict trials. VP and BLA contralateral inactivation did not affect the latency to press on the same side of the alley (inset). **E.** VP and BLA contralateral inactivations did not affect retrieval of auditory threat conditioning memory as shown by similar freezing levels in the inactivation group (n=6) as compared to the vehicle group (n=7). **F.** VP and BLA contralateral inactivation did not affect anxiety-like behavior and general locomotion in the open-field test, nor did it affect feeding behavior in the free food intake test, as indicated by similar time in the center of the open field and similar levels in the other behavioral measurements comparing vehicle (n=5) and inactivation (n=7) groups. *p<0.05, **p<0.01, and ***p<0.001 in post hoc comparison after ANOVA.

**Figure 5.**
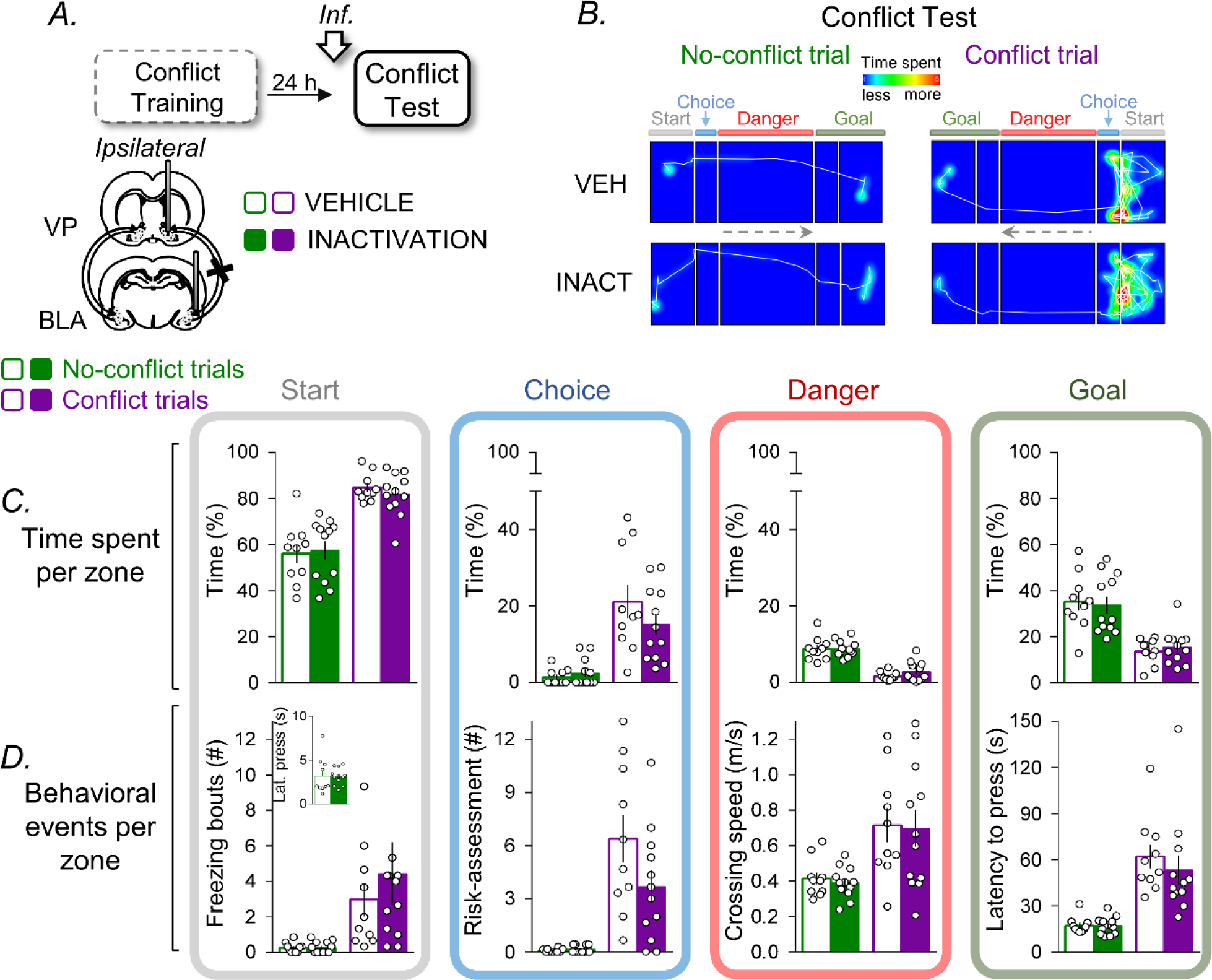
Ipsilateral VP-BLA inactivation did not affect crossing-related behaviors. To allow VP-BLA communication, rats were infused into the VP and BLA on the ipsilateral side with vehicle saline solution (VEH, n=10) or muscimol into VP and a cocktail of muscimol and baclofen into BLA (INACT, n=12) before the conflict test. **B.** Overlaid movement tracking and heat maps of representative rats (VEH and INACT) during the first two consecutive non-conflict and conflict trials of the conflict test. **C.** Analysis of time spent per zone showed that ipsilateral inactivation of the VP-BLA did not affect the crossing-mediated time spent in any of the zones. **D.** Analysis of behavioral events per zone shows that VP and BLA ipsilateral inactivation did not affect crossing-mediated behaviors, as well as latency to press on the same side of the alley (inset).

Consistent with our hypothesis, we observed that contralateral, but not ipsilateral, inactivation of the VP-BLA before the conflict memory test impaired the time spent and evaluated behaviors in most zones during conflict trials, while it left intact timing and behaviors triggered by non-conflict situations (**Figure 4A-B**). Similar to BLA inactivation, VP-BLA contralateral inactivation reduced the time spent in the start zone while increasing the time spent in the goal and danger zones (start zone, factorial ANOVA, group: F(1,26)= 6.34, p= 0.01; trials: F(1,26)= 62.90, p= 0.000; interaction: F(1,26)= 4.07, p= 0.053; conflict trials VEH: 86.0%, conflict trials INACT: 69.64%, post hoc comparison p= 0.017; goal zone, factorial ANOVA, group: F(1,26)= 6.78, p= 0.01; trials: F(1,26)= 52.53, p= 0.000; interaction: F(1,26)= 2.60, p= 0.11; conflict trials VEH: 12.65%, conflict trials INACT: 25.38%, post hoc comparison p= 0.02; danger zone, factorial ANOVA, group: F(1,26)= 1.96, p= 0.17; trials: F(1,26)= 59.92, p= 0.000; interaction: F(1,26)= 7.73 p= 0.009; conflict trials VEH: 1.34%, conflict trials INACT: 4.96%, post hoc comparison p= 0.03) (**Figure 4C**). Unlike BLA inactivation, however, contralateral inactivation did not affect freezing in the start zone (freezing bouts, factorial ANOVA, group: F(1,26)= 3.71, p= 0.06; trials: F(1,26)= 36.13, p= 0.000; interaction: F(1,26)= 3.88, p= 0.06), yet it slowed down crossing speed in the danger zone, thus delaying the latency to press in the goal zone (crossing speed, factorial ANOVA, group: F(1,26)= 2.10, p= 0.15; trials: F(1,26)= 15.31, p= 0.000; interaction: F(1,26) = 11.63, p= 0.002; conflict trials VEH: 0.76 m/s, conflict trials INACT: 0.49 m/s, post hoc comparison p= 0.01; latency to press, factorial ANOVA, group: F(1,26)= 28.84, p= 0.000; trials: F(1,26)= 67.02, p= 0.000; interaction: F(1,26)= 17.0, p= 0.000; conflict trials VEH: 54.58 s, conflict trials INACT: 25.61 s, post hoc comparison p= 0.000). Notably, similar to both VP and BLA inactivation, VP-BLA contralateral inactivation robustly impaired risk-assessment behavior in the choice zone (factorial ANOVA, group: F(1,26)= 26.83, p= 0.000; trials: F(1,26)= 65.05, p= 0.000; interaction: F(1,26)= 27.08, p= 0.000; conflict trials VEH: 7.12 events, conflict trials INACT: 1.61events, post hoc comparison, p= 0.000) (**Figure 4D**).

Next, we conducted separate experiments in different rat groups to explore whether contralateral VP-BLA inactivation was necessary for threat-related responses and feeding behavior. Unlike BLA inactivation, yet similar to VP inactivation, VP-BLA contralateral inactivation did not affect threat memory retrieval (VEH: 82.19%, INACT: 72.04%, Student’s two-tailed unpaired *t*-test, t(11)= −1.08, p= 0.29) (**Figure 4E**), while also leaving anxiety-like behavior, general locomotion, and feeding behavior intact (anxiety-like behavior, VEH: 1.50 s, INACT: 2.55 s, Student’s two-tailed unpaired *t*-test, t(10)= −0.64, p= 0.53; distance traveled, VEH: 29.71 m, INACT: 28.20 m, Student’s two-tailed unpaired *t*-test, t(10)= 0.71, p= 0.49; food-intake, VEH: 95.30%, INACT: 90.21%, Student’s two-tailed unpaired *t*-test, t(10)= 0.79, p= 0.44) (**Figure 4F**). The lack of an inactivation effect on the retrieval of threat memory was consistent with the lack of an inactivation effect on freezing bouts in the start zone during the CMC test.

In contrast to contralateral VP-BLA inactivation, ipsilateral inactivation during the conflict test had no effect on any of the temporal and spatial behavioral parameters evaluated (start zone, factorial ANOVA, group: F(1,40)= 0.04, p= 0.83; trials: F(1,40)= 63.22, p= 0.000; interaction: F(1,40)= 0.42, p= 0.5; choice zone, factorial ANOVA, group: F(1,40)= 0.89, p= 0.35; trials: F(1,40)= 39.31, p = 0.000; interaction: F(1,40) = 1.85, p = 0.18; danger zone, factorial ANOVA, group: F(1,40)= 0.76, p= 0.38; trials: F(1,40)= 89.72, p= 0.000; interaction: F(1,40)= 0.66, p= 0.41; goal zone, factorial ANOVA, group: F(1,40)= 0.00, p= 0.97; trials: F(1,40)= 42.32, p= 0.000; interaction: F(1,40)= 0.27, p = 0.6; freezing bouts, factorial ANOVA, group: F(1,40)= 0.50, p= 0.48; trials: F(1,40)= 10.57, p= 0.002; interaction: F(1,40)= 0.47, p= 0.49, risk assessment, factorial ANOVA, group: F(1,40) = 2.72, p= 0.10; trials: F(1,40)= 38.89, p = 0.000; interaction: F(1,40)= 3.01, p= 0.09; crossing speed, factorial ANOVA, group: F(1,40)= 0.02, p= 0.86; trials: F(1,40)= 14.74, p= 0.000; interaction: F(1,40)= 0.01, p= 0.90; latency to press, factorial ANOVA, group: F(1,40)= 0.48, p= 0.48; trials: F(1,40)= 41.03, p= 0.000; interaction: F(1,40)= 0.44, p= 0.50; latency to press in the start zone, Student’s two-tailed unpaired *t* test, t(6)= 0.08, p= 0.93) (**Figure 5A-D**). Thus, unlike the inactivation of the VP and BLA in the opposite hemispheres (contralateral), which had robust observable effects on various temporal and spatial behavioral parameters during the conflict test, inactivation in the same hemisphere (ipsilateral) did not result in any discernible impact on the parameters assessed. The specific and selective effect of contralateral inactivation suggests that the intricate interplay between VP and BLA is essential for orchestrating conflict-mediated behaviors, highlighting the nuanced and complementary roles of these structures in the regulation of risky actions.

In the CMC test, assessing risks in the choice zone emerged as a prominent behavior during conflict. Our inactivation experiments (including BLA, VP, and VP-BLA contralateral inactivation) significantly impaired this threat-related choice behavior, except for the ipsilateral inactivation experiment. Consequently, we conducted a separate analysis of this behavior to compare the experiments and evaluate the magnitude of the effects across the manipulations (**Figure 6A**). By normalizing the number of risk assessment events between trials relative to the control group, we revealed the specific impact of contralateral VP-BLA inactivation on risk assessment behavior. Notably, VP-BLA disconnection resulted in a more significant reduction in risk assessment compared to VP alone and BLA separately (BLA, non-conflict trials: z= −0.078, p= 0.93, conflict trials: z= - 1.075, p= 0.28; VP, non-conflict trials: z= 0.50, p= 0.61, conflict trials; z= −1.56, p= 0.11; contralateral, non-conflict trials: z= 0.08, p= 0.93; conflict trials: z= −2.18, p = 0.02; ipsilateral, non-conflict trials: z= 0.63, p= 0.52, conflict trials: z= −0.63, p= 0.52). These findings indicate that VP-BLA connectivity plays a vital role in controlling cautious approach choice behaviors guided by previous experiences (**Figure 6B**).

**Figure 6.**
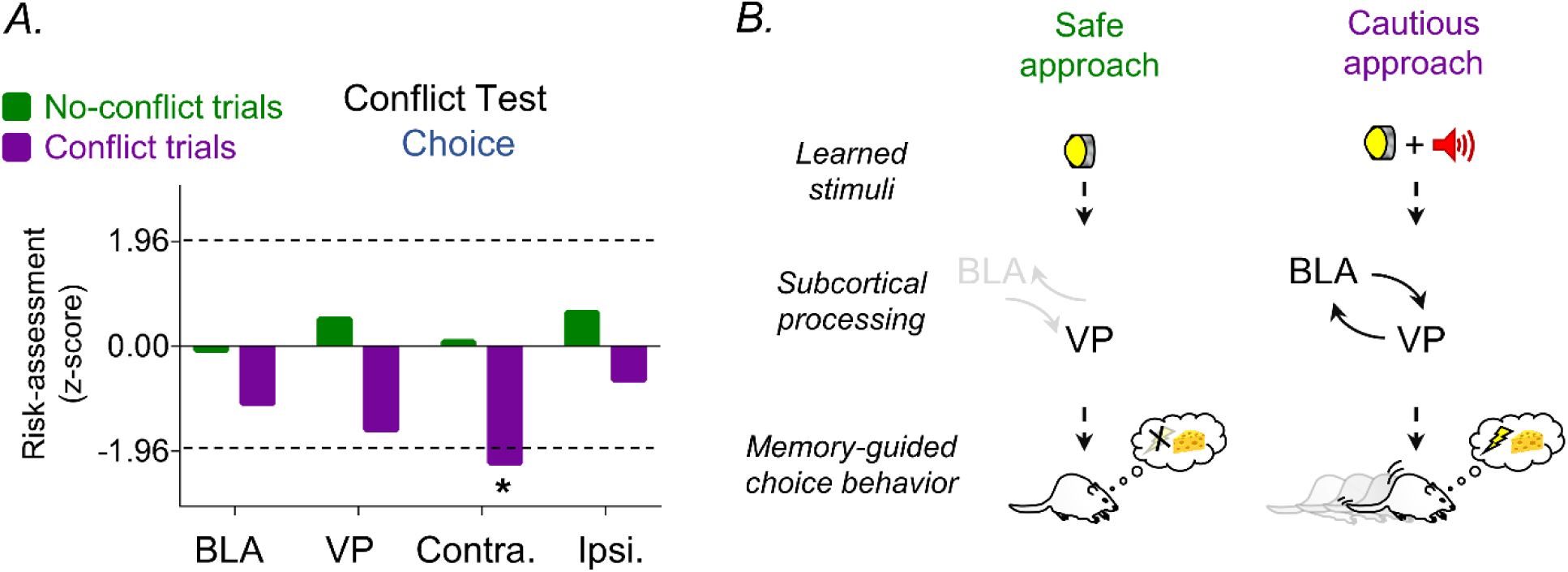
VP-BLA disconnection impairs conflict-induced risk taking. **A.** Data are normalized contrast of the number of risk assessments in the choice zone during the conflict test between trial types (non-conflict and conflict) across all inactivation experiments (BLA, VP, contralateral BLA-VP, and ipsilateral BLA-VP inactivations) relative to their respective control groups (z-score). Z-scores of risk assessments revealed that disconnecting the VP-BLA pathway significantly reduced (negative z-scores that exceeded −1.96; *p < 0.05) the drive to take risks in response to food-seeking conflicts. **B.** This diagram illustrates the proposed subcortical circuit for the cross-mediated conflict test. This highlights the key role of the VP in non-conflict trials, where efforts for food retrieval are prioritized (safe approach for food), and contrasts it with the involvement of VP-BLA connectivity in conflict trials, where cautious food-seeking persists despite associated threats (risky approach for food).

## Discussion

We investigated the individual and collaborative necessity of VP and BLA to influence foraging decisions in conflict situations. Our novel foraging choice task, coupled with pharmacological inactivation, revealed a new functional role for VP-BLA connectivity. Our findings align with previous evidence, consistent with the notion that BLA influences calculated choices toward threat-related behaviors, whereas VP invigorates choices toward reward-seeking responses. Importantly, our findings suggest that VP and BLA establish reciprocal feedback on the current value of threats and rewards, facilitating efficient decision-making during conflicts and appropriately restraining risky reward approach behaviors.

### Calculated Risk-Taking and Fear Suppression during Conflicts

Our study found that rats’ reaction times, patterns, and strategies for crossing to reach food changed depending on the presence or absence of threats, suggesting that rats make decisions that involve risk-taking, influenced by their food-seeking behavior. Previous research has shown that animals adjust their risk preferences based on the reward value, availability, threat intensity, and type. For example, rats exhibit increased risk-taking behavior in response to food deprivation, social competition, and predator cues (Davis et al., 2009; Choi and Kim, 2010; Dent et al., 2014; Engelke et al., 2021; Zoratto et al., 2022). The CMC task provides a realistic and ecologically relevant scenario for studying risk-taking behavior in rodents, involving calculated effortful actions that require suppressing defensive responses to threats (Illescas-Huerta et al., 2021). Prior research has shown that administering diazepam, a commonly used anxiolytic, before the CMC test encourages rats to cross more readily in pursuit of food when potential danger is involved, without impacting non-conflict situations (Illescas-Huerta et al., 2021).

This task is suitable for exploring active fear suppression during food-related conflicts, which differs from the passive fear suppression observed during fear extinction (Bouton, 2002; Sotres-Bayon and Quirk, 2010). We showed that VP-BLA connectivity is essential for risky reward behaviors involving active fear suppression. However, the role of VP in fear extinction remains unexplored. Comparing these fear suppression mechanisms (passive vs. active) may provide insights into how animals cope with various types of threats and balance their risks and benefits.

### Distinct Roles of BLA and VP in Foraging Choices

Our study uncovered distinct roles of the BLA and VP in shaping risk-taking behavior during food-seeking conflicts. Our results emphasize the central role of the BLA in shaping foraging decisions during conflicts, as it orchestrates delayed food-seeking decisions, promotes rapid crossing, and signals urgency in response to potential threats for food. These findings indicate that BLA plays a necessary role in food-seeking decisions that involve taking risks during conflicts, as it mediates the regulation of defensive responses to threats and likely gates of food-seeking behavior. This multifaceted involvement aligns with the role of BLAs in threat and reward processing as well as their ability to modulate approach-avoidance conflicts (Muller et al., 1997; Shabel and Janak, 2009; Choi and Kim, 2010; Sierra-Mercado et al., 2011; Amir et al., 2015; Kim et al., 2016; Piantadosi et al., 2017; Bindi et al., 2018). BLA’s role in encoding the value and salience of both aversive and appetitive stimuli, regulating emotional and motivational aspects of learning and memory, and its interaction with other brain regions such as prefrontal cortex, nucleus accumbens, and ventral tegmental area, highlight its importance in mediating the integration of threat and reward information and regulating goal-directed actions (Ambroggi et al., 2008; de Oliveira et al., 2011; Stuber et al., 2011; Sotres-Bayon et al., 2012; Namburi et al., 2015; Ramirez et al., 2015; Burgos-Robles et al., 2017; Diehl et al., 2020).

Our results indicate that VP plays a critical role in food-seeking behavior, particularly in weighing potential risks and rewards. The VP may be a key structure for survival decision-making, as it encodes the value and salience of appetitive and aversive stimuli and regulates decisions related to the motivational drive to seek food (Tindell et al., 2004; Richard et al., 2016; Saga et al., 2017; Richard et al., 2018; Moaddab et al., 2021). VP modulates the willingness to exert effort to obtain rewards and influences the selection of high-effort, high-reward options over low-effort, low-reward options (Farrell et al., 2021). VP mediates the effects of motivational manipulations, such as hunger and stress, on effort-related choice behavior (Farrar et al., 2008; Chang and Grace, 2014; Ottenheimer et al., 2020a). VP has been shown to increase the rate and intensity of operant responses such as lever pressing or nose poking, which are contingent on reward delivery (Farrar et al., 2008; Ahrens et al., 2018). VP also mediates the effects of reward anticipation, uncertainty, and magnitude on the vigor of actions (Ottenheimer et al., 2020b; Lederman et al., 2021; Moaddab et al., 2021). VP has been shown to interact with other brain regions, such as the lateral habenula, prefrontal cortex, and nucleus accumbens, to mediate the integration of reward and effort information, and the regulation of goal-directed actions (Mingote et al., 2008; Smith et al., 2011; Leung and Balleine, 2013; Tooley et al., 2018; Stephenson-Jones et al., 2020). The VP may be involved in determining the optimal level of effort required for food-seeking in the face of potential threats, likely via BLA connectivity.

### Collaborative Necessity of VP and BLA in Foraging Choices

VP and BLA have extensive mutual connections and may interact dynamically to balance the costs and benefits of risk-taking behavior and to adjust the motivational drive and vigor for food according to the environmental context (Zaborszky et al., 1984; Mitrovic and Napier, 1998; Mascagni and McDonald, 2009). We found that disconnecting the pathway between VP and BLA impaired the process of assessing and weighing the potential risks and rewards associated with food-seeking choices. This suggests that the connectivity between the VP and BLA plays a crucial role in controlling risky food-seeking choices during conflicts, while maintaining the motivation for food-seeking in non-conflict situations. Specifically, disrupting the VP-BLA pathway selectively impairs risk-taking behavior in response to food-seeking conflicts, highlighting its importance in assessing and weighing potential risks and rewards. This suggests that VP-BLA connectivity may be required to coordinate conflict-mediated behaviors, reconcile the interplay between reward and threat processing, and regulate risk-taking behaviors. This aligns with previous studies that have implicated both VP and BLA in threat and reward processing as well as in modulating approach-avoidance conflicts. In addition, our study reveals new perspectives on the connection between VP and BLA in the control of risky behaviors, which merits additional exploration. These results indicate that the inactivation effects of the contralateral VP-BLA are more similar to those of the BLA than the VP, implying that the BLA may have a more prominent role than the VP in mediating risky actions. This further highlights the importance of BLA-to-VP connectivity as a key neural pathway in survival decision-making, allowing animals to restrain risky behaviors and adopt a cautious approach when seeking food.

### Caveats and future directions

Our study has several limitations that are currently being addressed in ongoing experiments. First, our research found that rats displayed varying levels of risk-taking behavior, even within the same group. Future studies should investigate the origins and consequences of these individual differences, and how they may influence neural and behavioral responses. Second, our study only involved male rats; therefore, future research should examine female rats (Orsini et al., 2016). Third, our method of pharmacological inactivation did not differentiate between specific cell types or pathways. Future research should employ more refined techniques, such as optogenetic manipulation and single-unit recordings, to gain a deeper understanding of the specific cell and pathway mechanisms that govern neural processes that promote adaptive behaviors. In particular, it would be intriguing to investigate the unique contributions of specific cell types in the VP and BLA during conflict situations, as certain neuronal types in both of these regions have been linked to approach or avoidance behaviors (Beyeler et al., 2018; Stephenson-Jones et al., 2020).

## Conclusions

Our research uncovered a new function of the VP-BLA connection in regulating foraging decisions in situations of conflict. We demonstrate that VP and BLA work together to assess and signal threats and reward responses and that their communication is essential for balancing approach and avoidance behaviors. Our findings provide novel insights into the neural mechanisms that govern survival decision-making in animals and suggest that targeting the VP-BLA connection may offer potential for treating disorders related to impaired risk-taking behavior, such as addiction, anxiety, and depression.

## Methods

A total of 181 adult male Wistar rats (Institute of Cell Physiology breeding colony), aged 2-3 months and weighing 250-300 g, were individually housed in polyethylene cages. Daily handling aimed at minimizing stress responses, and the rats adhered to a standard 12-hour light/dark schedule. Ten rats were excluded from the data analysis due to CMC task learning difficulties, while thirty-four were excluded based on histological findings or insufficient post-surgery recovery. All the experiments were performed during the light phase. To maintain consistent motivation for food-related tasks, the rats underwent food restriction (12 g/day of standard laboratory rat chow, with a 5 g bonus feeding weekly to sustain rats at 85% of their initial weight). The procedures were approved by the Institutional Animal Care and Use Committee of the Universidad Nacional Autónoma de México following the guidelines of the National Ministry of Health for Laboratory Animal Care.

### Crossing-Mediated Conflict task

The task involved training and testing in straight alleys (100 × 30 × 50 cm) with stainless-steel frames situated in a sound-attenuating chamber (150 × 70 × 140 cm). These alleys comprise two safety zones and one danger zone (**Figure 1A**). The danger zone (60 cm × 30 cm) had a stainless-steel bar floor capable of delivering a scramble footshock (Coulbourn Instruments, USA). The safety zones (20 cm × 30 cm) included a speaker, cue light, lever linked to a pellet dispenser, food plate, and acrylic floor. A computer with custom scripts managed pellet delivery, cue lights, speakers, shock delivery, and recorded lever presses. Between the experiments, the alley’s floors and walls were cleaned with soapy water and 70% alcohol and dried with paper towels. Before conflict training, all rats were conditioned to associate lever pressing with sucrose pellet reinforcement (45 mg dustless precision pellets; Bioserve, USA). Each session began and concluded with a 5-minute context-alone exposure devoid of cue lights, shocks, or noise.

#### Conflict training

Rats underwent training for the crossing-mediated conflict task as previously described (Illescas-Huerta et al., 2021). Briefly, this training encompassed three stages: reward training, threat training, and discrimination training to differentiate between non-conflict and conflict trials. The entire conflict training spanned 34 days, with a single daily training session.

##### Reward training

Rats confined to a safety zone were trained to anticipate food availability with the presentation of a light cue. When the cue light was illuminated, a pellet was delivered to the food plate with each lever pressed (light trial); however, no pellet was delivered in the absence of light (no-light trial). Both light and no-light trials were randomly presented per session. After rats showed an increased and stable lever press during light trials, but not during no-light trials, they were trained to cross from one safety zone to the other safety zone to the other to obtain food cued by light. A reward-crossing trial started when the light turned on, on the side of the alley opposite the location of the rat. The trial ended when the light turned off after either the rat had crossed to that safety zone and pressed the lever, or after 180 seconds without crossing. Rats received one to three reinforced lever presses on the same safety zone to prevent crosses performed as non-signaled habitual reactions. Up to this point, rats were required to acquire food by crossing to the safety zone in the alley where the light was illuminated and pressing the lever, without encountering conflict.

##### Threat training

Rats confined to the danger zone, experienced five pairings per session (variable inter-trial interval of 1-3 minutes) of white noise (85 dB, 30 s) coupled with a mild foot shock (0.5 mA, 1 s). The shock occurred randomly during the tone to avoid anticipation of delivery time. The next day, two noise-alone trials in the absence of shocks were presented to test cued threat memory. Next, rats were challenged to cross the alley in the same way as in reward crossing trials, but in the presence of a threat consisting of noise signaling, a foot shock of 0.5 mA in the middle of the alley (variable inter-trial intervals ranged from 1 to 3 min). Each trial ended either after the time of crossing had elapsed (30-120 s) or when the rats successfully crossed and pressed the lever on the opposite side of the alley. By this point, the rats had learned to cross to obtain food despite threats indicated by the concurrent light and foot shock signaled by sound cues during the conflict.

##### Discrimination training

Rats were trained to discriminate between non-conflict (crossing for food without threat) and conflict (crossing for food despite threat) trials. Short acrylic barriers (9 cm tall) were strategically placed between each safety zone and the grid, defining a choice zone to constrain choice-mediated crossing behavior in both time and space. Each discrimination session comprised three consecutive blocks of ten crossing trials, with seven or nine non-conflict trials, and one or three conflict trials per block (10% and 30% chance of threat trials, respectively). The trial types were presented randomly to prevent anticipation. The rats had limited time (180 s) to decide whether to cross the alley in each trial. The proportion of conflict trials and shock intensity increased during the training period. After this stage, the rats underwent cannula-implantation surgery. Upon full recovery, they underwent four additional discrimination sessions, each with three blocks containing seven non-conflict and three conflict trials. Shock intensity increased by 0.1 mA per day, reaching 0.8 mA. Rats failing to discriminate trial types by the end of training (p > 0.05) on the last day between conflict and non-conflict trials and those unable to cross within 180 s (10 of 108 rats) were excluded from the study.

#### Conflict test

The conflict test was conducted on the day following the additional discrimination sessions. Rats underwent ten crossing trials presented under the same conditions as the last day of additional discrimination training (30% chance of threat: three conflict and seven non-conflict randomly presented trials), but without shocks. Results were expressed by contrasting the average temporal and spatial behavioral patterns, encompassing cues, crossing, reward-seeking actions, and threat responses, between the control and experimental groups during both conflict and non-conflict trials.

### Other behavioral tests

#### Threat memory test

Training and testing were conducted using conditioning chambers (Coulbourn Instruments). On the first day, threat conditioning involved five trials with a tone (75 dB, 30 s) co-terminated with a foot shock (0.7 mA, 1 s) delivered through a stainless-steel grid floor. On the second day, the threat memory test consisted of two trials with tone presentations in the absence of a foot shock. Trials started after 5 min without stimuli and followed a variable inter-trial interval of 120 s.

#### Free food intake test

The test was conducted in standard home cages, where rats were provided with a 20 g plate of Bioserve (45 mg, dustless precision pellets) food for 30 min.

#### Open field test

The test involved individually placing the rats in an arena with a dimly lit (20 lx) and recording their behavior for 10 min. Locomotion was assessed based on the distance traveled, while anxiety-like behavior was evaluated based on the center time in the arena.

### Stereotaxic surgery

Rats were anesthetized with isoflurane inhalant gas and positioned in a stereotaxic apparatus. Anesthesia maintenance (2-3% isoflurane) was facilitated using a facemask. For pharmacological inactivation, bilateral implantation of 23-gauge stainless steel guide cannulas (15 mm hypodermic stainless steel; AM Systems) occurred in the BLA (−2.8 mm AP; ± 5.0 mm ML; −7.5 mm DV, from Bregma) and VP (+0.4 mm, AP; ±2.1 mm, −7.5 DV, from Bregma). In the disconnection experiments, guide cannulas were placed contralaterally or ipsilaterally in the BLA and VP, with the hemisphere side counterbalanced. Cannula tips were aimed at 1.0 mm above the target structure, secured with dental acrylic cement, and anchored with two surgical screws on the skull. Rats were allowed a minimum of seven days for post-surgery recovery before initiating behavioral training.

### Local infusions

The day before infusion, the rats were habituated to manipulation to minimize stress, with injectors briefly inserted into the cannulas without infusion. The injector tips were extended by 1 mm beyond the guide cannula. On the test day for each behavioral task, muscimol and baclofen (Sigma-Aldrich) GABA receptor agonists, or muscimol alone were used to inactivate target regions (INACT). Muscimol and baclofen (125 ng of each drug/0.5µl per side; as in: Ghods-Sharifi et al. (2009)) or vehicle saline solution (VEH) was infused into BLA. Muscimol (10 ng/0.5µl per side, as in: Farrar et al. (2008)), or VEH was infused into the VP. Infusions were performed 15 min before behavioral testing. The rate of infusion was 0.4 µl/min controlled by a micro infusion pump (KD Scientific) operating ten µl syringes (Hamilton) connected to the cannulas via polyethylene tubing. The injectors were left in place for 1 min after infusion to allow diffusion.

### Histology

After the behavioral experiments, the rats were deeply anesthetized with pentobarbital sodium (150 mg/kg, i.p.) and transcranially perfused with 0.9% saline. Brains were stored in 10% formalin solution (Sigma-Aldrich) for at least three days. Formalin was then substituted with 30% sucrose solution until tissue saturation was reached. Brains were cut into 40-μm-thick coronal sections using a cryostat (Leica CM1520), stained for Nissl bodies, and examined under a bright-field microscope (Nikon, H550S) to verify cannula tip placement. Only rats with cannula locations within the borders of the BLA and VP were included in statistical analysis.

### Data collection and analysis

All behaviors were recorded using digital video cameras (Provision, model D-380D5) located above each task apparatus. Conflict test video images were analyzed using the video tracking software ANY-maze 7.1 (Stoelting, USA). Temporal and spatial behavioral patterns, including cues, crossing, reward-seeking actions, and responses to potential threats, were assessed by segmenting the 100 cm alley into four tracking zones: start (19 cm), choice (10 cm), danger (42 cm), and goal (29 cm). The start zone marked the safety zone where rats initiated crossing trials, whereas the choice zone represented the area for risk assessment before crossing, encompassing acrylic barriers and the initial three stainless steel bars of the grid. The danger zone comprised the grid where the rats crossed, and the goal zone denoted the safety zone where the rats concluded the crossing trial by pressing a lever. Rat tracking at three body points (head, center, and tail) provided data on the time spent and entries per zone. The center point of the rat was used to determine its presence in the start, danger, and goal zones, whereas the head point was used to assess its presence within the choice zone (with the head point required to be within 10 mm of the zone for a successful visit). The percentage of time spent per zone was calculated as follows: (time spent per zone × 100) / (total time spent in the start, danger, and goal zones). Additionally, freezing bouts (number of times the rats spent at least 300 ms immobile in the start zone), latency to press (seconds to press a lever on the same side), risk assessment events (number of entries into the choice zone), and crossing speed (rate (m/s) at which the rat crosses the danger zone) were analyzed. Finally, we calculated the latency to press in the goal zone, which represented the combined time spent in the start, danger, and goal zones. This measurement determined the reaction time required to complete the crossing from the start zone to the goal zone.

Movement tracking and heat maps were obtained from the rat head position. A maximum of 10 s was used as the hottest color in the heat map. All data, movement tracking, and heat maps were obtained for both the conflict and non-conflict crossing trials. Freezing responses were expressed as the percentage of time spent without movement (300 ms immobile, except for respiration) during the auditory cue presentation. The percentage of freezing, distance traveled (m), and center time in the open field were calculated automatically using tracking software (ANY-maze 7.1, Stoelting, United States). Food intake was obtained by comparing the food plate weight (g) before and 30 min after food presentation. The percentage of food intake was calculated as follows: (total food intake × 100) / (total amount of food presented). Food intake was manually recorded and scored by an experimenter blinded to the experimental conditions. Experimental groups were compared by using, when appropriate, unpaired Student’s two-tailed *t*-tests or analysis of variance (two-way ANOVA) followed by planned comparisons or Tukey’s multiple comparisons post hoc test (STATISTICA; Stat Soft, USA and GraphPad Prism 7). The accepted value for significance was set at p < 0.05.

## Acknowledgments

This study was supported by the Consejo Nacional de Ciencia y Tecnología (CONACyT, grant PN2463), as well as by the Dirección General de Asuntos del Personal Académico de la Universidad Nacional Autónoma de México (UNAM, grants IN205417 and IN214520) and the International Brain Research Organization (Return Home fellowship) to FS-B. A H-J is a doctoral student at Programa de Doctorado en Ciencias Biomédicas at UNAM and was supported by a CONACyT fellowship (417151). We thank Dr. Christian Bravo-Rivera for his comments on the manuscript and the Sotres-Bayon laboratory members for their technical assistance and helpful discussions.

